# A population-based temporal logic gate for timing and recording chemical events

**DOI:** 10.1101/029967

**Authors:** Victoria Hsiao, Yutaka Hori, Paul W.K. Rothemund, Richard M. Murray

## Abstract

Single-cell bacterial sensors have numerous applications in human health monitoring, environmental chemical detection, and materials biosynthesis. Such bacterial devices need not only the capability to differentiate between combinations of inputs, but also the ability to process signal timing and duration. In this work, we present a two-input temporal logic gate that can sense and record the order of the inputs, the timing between inputs, and the duration of input pulses. The temporal logic gate design relies on unidirectional DNA recombination with bacteriophage integrases to detect and encode sequences of input events. When implemented in a chromosomally-modified *E. coli* strain, we can utilize stochastic single cell responses to predict overall heterogeneous population behavior. We show that a stochastic model can be used to predict final population distributions of this *E. coli* strain, and thus that final differentiated sub-populations can be used to deduce the timing and duration of transient chemical events.

---

Subject categories: Synthetic biology, DNA memory, Modeling biology

## Introduction

Engineered bacteria could one day be powerful self-replicating single-cell sensors with environmental, health, and industrial applications. Synthetic biology has made important strides in identifying and optimizing genetic components for building such devices. In particular, much work has focused on Boolean logic gates which detect the presence or absence of static chemical signals (Gardner *et al*, 2000; Anderson *et al*, 2007; Wang *et al*, 2011; Moon *et al*, 2013; Shis *et al*, 2014) and compute a digital response.

Temporal logic gates, which process time-varying chemical chemical signals, have been much less explored. Pioneering work by Friedland *et al*. used serine integrase-based recombination for the counting and detection of sequential pulses of inducers (Friedland *et al*, 2009). But so far, no work has studied the potential for temporal logic gates to provide information about the duration of a signal, or the time between two chemical events. Here, we present a temporal logic gate which allows us to infer analog signal timing and duration information about the sequential application of two inducer molecules to a population of bacterial cells.

Similar to previous temporal logic gates, our design takes advantage of the irreversibility of serine integrase recombination. While bistable switches have been successfully deployed as memory modules in genetic circuits (Kotula *et al*, 2014), such switches require constant protein production to maintain state, and are sensitive to cell division rates and growth phase. The large serine integrases, on the other hand, reliably and irreversibly flip or excise unique fragments of DNA (Yuan *et al*, 2008). Thus logic circuits built from integrases intrinsically include DNA-level memory that requires virtually no cellular resources to maintain state, thus enabling permanent and low-cost genetic differentiation of individual bacterial cells based on transient integrase induction. Further advantages of the serine integrates include the short length (40-50 bp) and directionality of their attachment sites. Serine integrases recognize flanking DNA binding domains (attB, attP) and subsequently digest, flip or excise, and re-ligate the DNA between the attachment sites. Flipping or excision activity is determined by the relative orientation of the sites, which allows complex orientation-dependent behavior to be programmed into integrase circuits. Well-known serine integrases include Bxb1, TP901-1, and *ϕ*C31, all of which have been used to demonstrate static-input logic gates (Siuti *et al*, 2013; Bonnet *et al*, 2013), and some have cofactors that can reverse directionality (Bonnet *et al*, 2012; Khaleel *et al*, 2011). Recently, an entirely new set of 11 orthogonal integrases was characterized, greatly expanding the set of circuits that can be built (Yang *et al*, 2014).

In contrast to previous studies of temporal logic gates, our work leverages the stochastic nature of single-cell switching to create a robust population-level response to a time-varying chemical signal. By traditional engineering standards, synthetic circuits would ideally perform identically in every cell in a population. When this ideal is applied to biology, the stochastic nature of molecular processes, particularly at low copy numbers, presents a significant barrier to reliable outputs from engineered cells. Thus while natural cellular dynamics and differentiation take advantage of noisy gene expression (Elowitz *et al*, 2002; Süel *et al*, 2007) synthetic circuits often require noise reduction for proper function (Dunlop *et al*, 2008). We designed a two-input temporal logic gate using strategically interleaved and oriented integrase (Bxb1, TP901-1) DNA recombination sites and used this gate to engineer an E. coli strain with four possible genetically-differentiated end states. This strain contained single genomic copies of the temporal logic gate, ensuring digital-yet-stochastic responses from individual cells. We then utilized the heterogeneity of individual cellular responses to encode sequences of chemical inputs into the overall population response, and use a stochastic model of single cell trajectories to predict the population response. By analyzing the distributions of final cell states, we deduce the timing and pulse duration of transient chemical pulses. Furthermore, because the states are genetically encoded, we can recover details of a chemical event long after its occurrence.

## Results

### Design of a two-integrase temporal logic gate

We have designed a two-input temporal logic gate that differentiates between the start times of two chemical inputs and produces unique outputs accordingly (Figure 1A). The design relies on a system of two-integrases with nested integrase attachments sites (Figure 1B). The use of integrases irreversibly invert segments of DNA, resulting in a memory feature that can be maintained for multiple generations (Bonnet *et al*, 2012).

**Figure 1:**
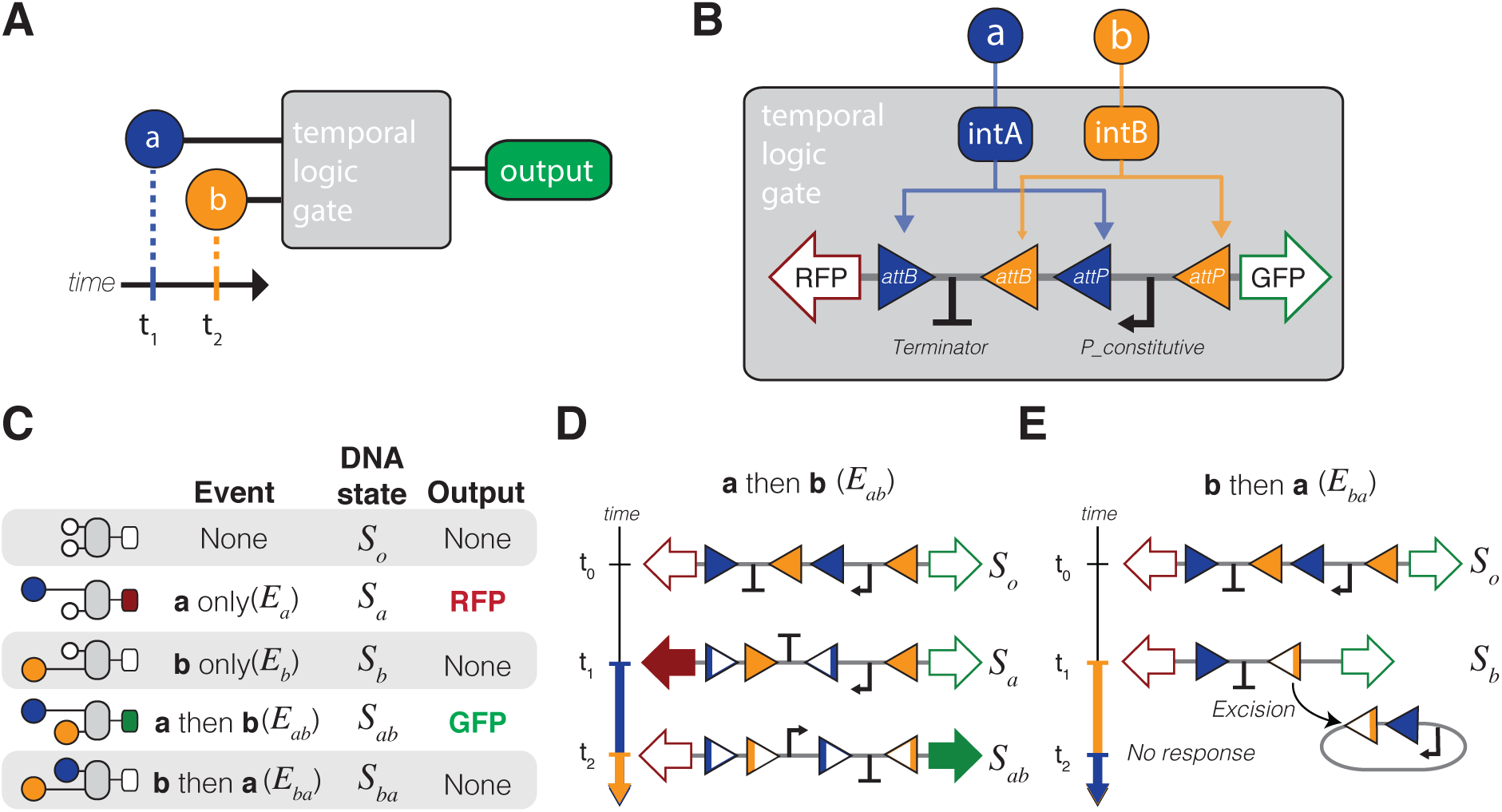
Design overview of a temporal logic gate. A) A temporal logic gate distinguishes between two chemical inputs (**a,b**) with different start times. B) Implementation of the temporal logic gate using a set of two integrases with overlapping attachment sites. Chemical inputs **a** and **b** activate production of integrases intA and intB, which act upon a chromosomal DNA cassette. C) Table with all possible inputs and outcomes to the event detector. D) Sequence of DNA flipping following inputs with inducer **a** before inducer **b** (event *E_ab_*). E) Sequence of DNA flipping following inducer inputs with **b** first (event *E_ba_*). In any events in which **b** precedes **a**, the uni-directionality of the intB attachment sites results in excision.

The design of the integrase temporal logic gate hinges on interleaving the attB attachment site of integrase B (intB) with the attP site of integrase A (intA), thus ensuring that the possible DNA flipping outcomes are mutually exclusive (Figure 1B). The serine integrases used in this design are TP901-1 (intA) and Bxb1 (intB). The fluorescent proteins mKate2-RFP (RFP) and superfolder-GFP (GFP) are used as placeholders for future downstream gene activation as well as real-time readouts of the logic gate. The design also features a terminator (Bba-B0015) and a strong constitutive promoter (P7). In the case where there are no inputs, the terminator prevents expression of RFP from the constitutive promoter.

There are five possible basic events that could occur in a two-input system (Figure 1C): no input, inducer **a** only (*E_a_*), inducer **b** only (*E_b_*), inducer **a** followed by **b** at a later time (*E_ab_*), and inducer **b** followed by **a** at a later time (*E_ba_*). Consequently, in a perfectly resolved temporal logic gate there should be five unique DNA states corresponding to the five types of events: *S_o_* (the initial state), *S_a_, S_b_, S_ab_*, and *S_ba_*. This design is limited to only four DNA states due to excision when *E_b_* occurs (*S_b_* = *S_ba_*). The two fluorescent outputs correspond to the two states that occur when inducer **a** is detected first – RFP is produced when the cell is in state *S_a_*, and GFP is produced when the cell is in state *S_ab_*.

Figure 1D illustrates the sequence of recombination that occurs during an event *E_ab_* that results in DNA state *S_ab_* and the production of GFP. Upon addition of inducer **a** at time *t*_1_, TP901-1 flips the DNA between its attachment sites, reversing the directionality of the terminator and the Bxb1 attB recognition site (state *S_a_*). Then, when inducer **b** is added at some time *t*_2_ that is greater than *t*_1_, the directionality of the Bxb1 sites is such that the DNA is flipped to reverse the directionality of the P7 constitutive promoter (state *S_ab_*). If inducer **b** is added first (Figure 1E), the Bxb1 attachment sites are uni-directional, a configuration that results not in recombination, but in excision of the DNA between the sites (state *S_b_*).

Once a DNA recombination has occurred, it is irreversible. The unique attB and attP attachment sites are recombined into attL and attR sites, respectively, and no longer recognized by the integrases. The nesting of the integrase attachment sites is the key design feature that produces the temporal ***a*** *then* ***b*** logic, and the irreversibility of the recombination records the event in DNA memory. The result is a genetic record that can both be sequenced later and immediately read by via constitutive production of fluorescent outputs.

### A Markov model for integrase recombination

We created a model of integrase-mediated DNA flipping and then used a stochastic simulation algorithm (Gillespie, 1977) to simulate individual cell trajectories (Figure 2A). All of the four possible DNA states are represented in the model: the original state (*S_o_*), the intB excision state (*S_b_*), the intA single flip state (*S_a_*), and the ***a*** *then* ***b*** double flip state (*S_ab_*). We have implemented the system experimentally by chromosomally integrating the target DNA into the genome of the *E. coli* cell. This allows us to assume that *ea*ch cell only has one copy of the temporal logic gate (Haldimann and Wanner, 2001), and that *ea*ch cell can be characterized by the tuple (DNA, IntA, IntB) (Figure 2B). The DNA term is *S_o_*, *S_a_, S_b_*, or *S_ab_*, and IntA and IntB are non-negative integers representing the molecular copy number of *ea*ch integrase. Once a DNA cassette has flipped into any of the states other than the original state *S_o_*, there is no reverse process. The logic gate is designed such that if integrase B is expressed prior to integrase A, the DNA cassette is excised and the chain reaches the dead-end *S_b_* state. In order for a cell to successfully detect *E_ab_*, it first needs to switch into state *S_a_* then transition into state *S_ab_* upon addition of inducer **b**.

**Figure 2:**
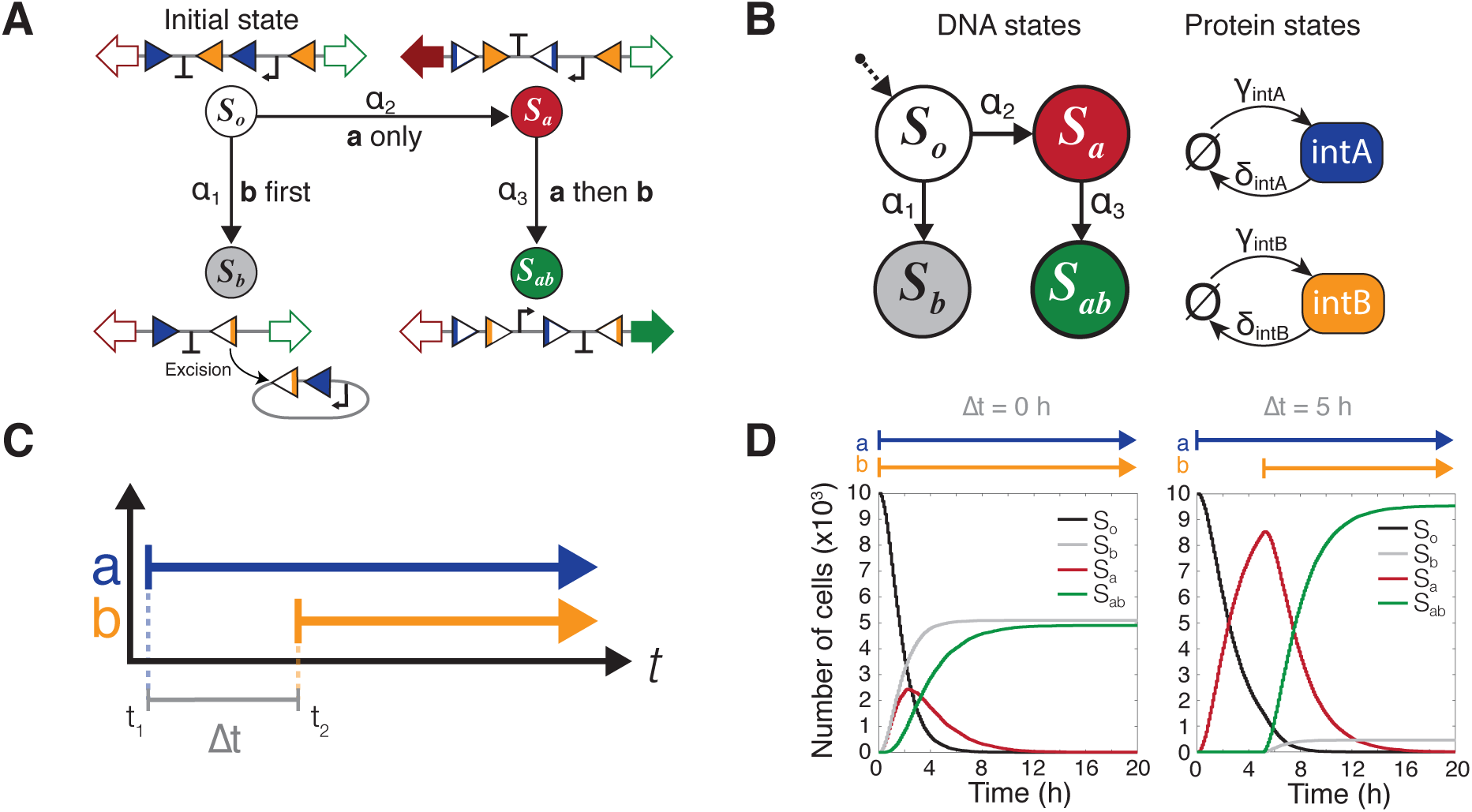
A Markov model of integrase-mediated DNA flipping. A) The four possible DNA states, illustrated with DNA state diagrams. All DNA begins in the initial state *S_o_* and there are no reverse processes. The propensity functions *α*_1_*,α*_2_, and *α*_3_ are dependent on the concentration of the two integrases and correspond to the events ***b*** *first* (*E_b_*), ***a*** *only* (*E_a_*), and ***a*** *then* ***b*** (*E_ab_)*, respectively. B) Representation of the same model as a Markov chain. Integrases are represented simply as protein states with production (*γ*_A_, *γ_b_*) and degradation (*δ_A_, δ_B_*) rates. C) Graphical representation of inducer step functions. Δt is defined as difference between the start time of the first inducer and start time of the second. D) Simulation results for inducer separation times of 0 and 5 hours. There are four possible DNA states, but all cells end up in either the *S_b_* or *S_ab_* final states. Individual trajectories are simulated for 10,000 cells and the number of cells in each DNA state are summed for each time point (Figure S1).

Since each cell contains only a single copy of the temporal logic gate DNA, we can expect each cell to behave differently, and to be highly susceptible to internal and external noise. This stochastic behavior will create a heterogeneous population response that can be analyzed for a more complex profile of event than if all the cells behaved uniformly. In order to capture the heterogeneity of cell population, we model the temporal logic gate using a stochastic model. Specifically, the stochastic transitions between the DNA states and the production/degradation of integrases are mathematically modeled by a continuous-time Markov chain over the state space (DNA, IntA, IntB) as illustrated in Figure 2B. Definitions of transition rates can be found in the Supplementary Information (Table S4).

*In silico*, the dynamics of a single cell translates to each stochastic simulation of the Markov model starting with (DNA = *S_o_*, IntA = 0, IntB = 0) state. We define ℙ*_t_*(*S_o_*), ℙ*_t_*(*S_a_*), ℙ*_t_*(*S_b_*) and ℙ*_t_*(*S_ab_*) as the probability that the DNA state of a single cell is *S_o_, S_a_, S_b_* and *S_ab_* at time *t*, respectively. The temporal dynamics of the probability can be modeled by the following ordinary differential equation (ODE) (see SI for derivation).

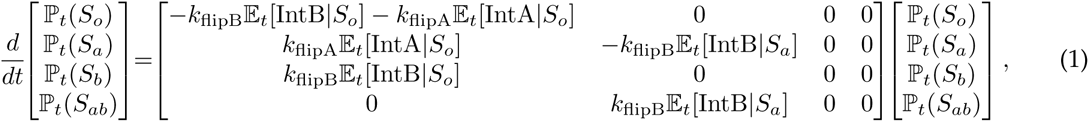
where the notation 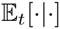 stands for the conditional expected value at time *t*.

We define the time between the introduction of the first inducer (*t*_1_) and the arrival of the second inducer (*t*_2_) as the *inducer separation time* (Δ*t*), such that

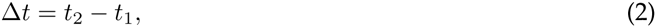
as shown in Figure 2C.

In the following set of simulations and experiments, we will consider cases with step inputs (Figure 2C), where the inducers are either present or not present. Concentrations of the inducers when they are “on” will be held constant. Also, it is important to note that inducer **a** is still present during and after time Δ*t* when inducer **b** is introduced.

Simulations of the Markov model were done with biologically plausible parameters in order to predict qualitative circuit behavior (Table S4). We limited the parameters to only the basic processes (integrase production, degradation, and DNA flipping), and parameter values were chosen to be within biological orders of magnitude. The single production rate constants, *k*_prodA_ and *k*_prodB_, combine the transcription and translation rates of each integrase. When an integrase in the model is induced, its production rate, γ*, is the sum of *k*_prod_* and any leaky transcriptional expression, *k*_leak_* (* = intA or intB). Any individual integrase, once produced, will need to dimerize, search for the DNA binding site, bind to the DNA, form a dimer of dimers (tetramer), digest, flip, and ligate the DNA (Yuan *et al*, 2008). We have combined all of those rates into the rate constant, *k*_flipA_ and *k*_flipB_. Parameter values for preliminary simulations were *k*_prodA_ = *k*_prodB_ = 0.5(*μ*m^3^∙hr)^−1^, *k*_deg_ = 0.01hr^−1^ (69 min half-life), *k*_flipA_ = *k*_flipB_ = 1hr^−1^, and *k*_leakA_ = *k*_leakB_ = 0(*μ*m^3^ ∙ hr)^−1^.

Our analysis of initial numerical simulation results underscored the significant role that the inducer separation time, Δ*t*, plays in setting the final population distributions (Figure 2D). For each Δ*t*, individual cell trajectories (N = 10,000) were generated with the assumption that each cell only has one copy of the target DNA. Then, at every time point, the total number of cells in each DNA state was counted (Figure S1). Figure 2D shows the contrast between adding both inducers simultaneously (Δ*t* = 0h) and adding inducer **b** after a 5 hour delay (Δ*t* = 5h). Since both inducers are present by the end of simulation, all of the cells must have a final state that is either the *S_ab_* state or the *S_b_* state. No cells remain in the original *S_o_* configuration. *S_a_* is a transient state that builds up prior to the addition of inducer **b** and begins to convert to *S_ab_* immediately after the introduction of **b**. These initial simulation results suggest that Δ*t* may be a way to reliably tune the final population fractions of *S_ab_* versus *S_b_* state cells.

### Characterization of inducer separation time

We can further investigate the effects of varying both inducer order and separation time *in silico* (Figure 3). Populations of cells have again been exposed to a sequence of overlapping step functions (N = 10,000). In the case of an *E_ab_* event, the proportion of cells that successfully detect *a then b* and switch to state *S_ab_* is a function of the inducer separation time, Δ*t* (Figure 3A). Exposing cells to the inverse sequence of events, *E_ba_*, results in a decrease of *S_ab_* cells proportional to increasing Δ*t* (Figure 3B). If we plot the final number of *S_ab_* cells from both *E_ab_* and *E_ba_* as a function of Δ*t* (Figure 3C), we see that the two curves do not overlap. This indicates that just the proportion of *S_ab_* cells alone is enough to uniquely distinguish which inducer appeared first. Simulation results tracing the other three DNA states can be found in Figure S2.

**Figure 3:**
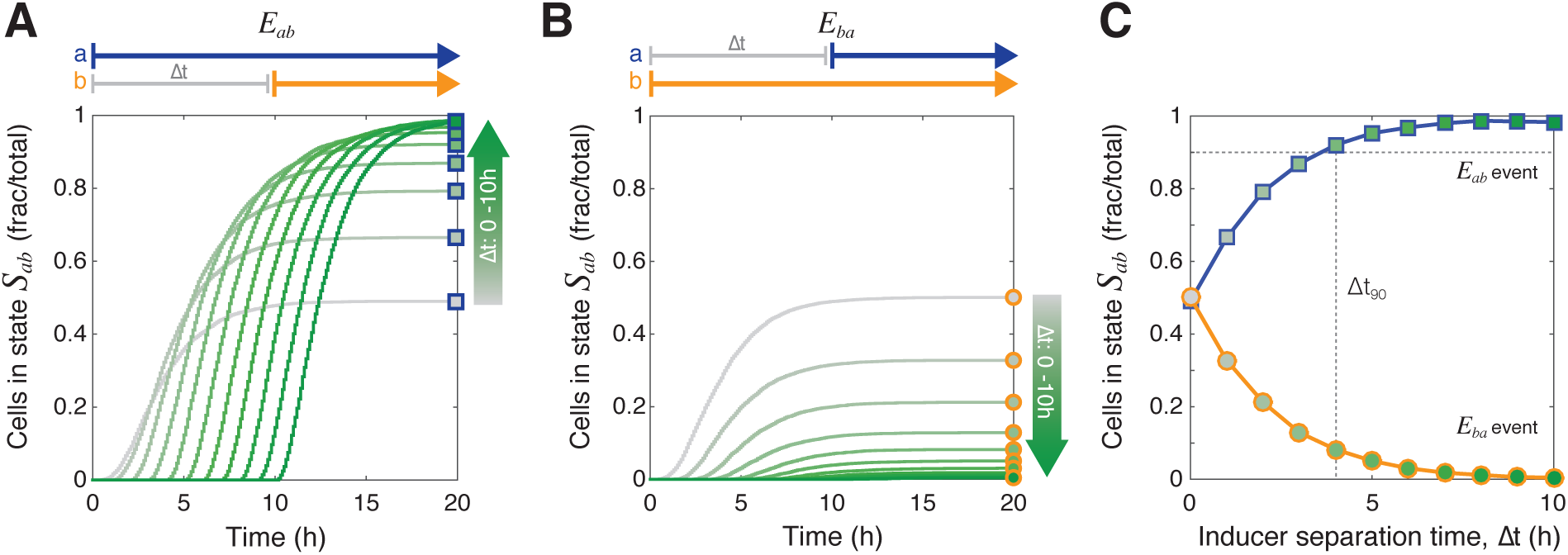
Simulation results for inducer separation time for Δ*t* = 0 − 10h. A) The population fraction (N/10,000 cells) that switches into state *S_ab_* following an *E_ab_* event is dependent on the inducer separation time, Δ*t*. The gray to dark green color gradient represents increasing Δ*t* values. Square markers indicate final population fractions for specific values of Δ*t*. B) In the case of the inverse *E_ba_* event, the fraction of cells in state *S_ab_* decreases proportionally to Δ*t*. Circular markers indicate final population fractions for specific values of Δ*t*. C) Final *S_ab_* cell fractions from Figure 3A, B are plotted as a function of Δ*t*. Blue line with square markers are endpoint population fractions from an *E_ab_* event. Yellow line with circular markers are final endpoint population fractions from an *E_ba_* event. The gradient inside the markers corresponds to increasing Δ*t* value. The dotted gray line corresponds to the Δ*t*_90_, the value of Δ*t* at which 90% of the cells are in state *S_ab_*. All simulations were done with a population of N = 10,000 cells.

Additionally, we can define a detection limit, Δ*t*_90_, for which the inducer separation time results in 90% of population switching into the *S_ab_* state (Figure 3C). This Δ*t*_90_ limit provides a way to capture the two response regimes of the population. If the inducer separation time is less than the detection limit (Δ*t* < Δ*t*_90_), then the rate of population switching is fast enough such that the number of *S_ab_* cells will correspond uniquely to some Δ*t* value. If Δ*t* > Δ*t*_90_, then most cells have already switched to a final state, and the differences in *S_ab_* cell count are too small to uniquely determine Δ*t*.

*In vivo* experimental results showed that population fractions of *S_ab_* cells could be tuned using Δ*t*, and verified model predictions (Figure 4). DH5*α*-Z1 cells were chromosomally integrated with one copy of the integrase target DNA and then transformed with a high copy plasmid containing Ptet-Bxb1 and PBAD-TP901-1. When Δ*t* is varied from 0 to 8 hours, we observed results qualitatively similar to model predictions (Figure 4). In Figure 4A, the cells have been exposed to an *E_ab_* event, where inducer **a** is present from time *t* = 0 h to *t*_end_, and **b** is present from *t* = Δ*t* h to *t*_end_. GFP expression is a proxy for *S_ab_* state cells, and total fluorescence has been normalized by a non-induced control sample. The number of cells in the GFP-expressing *S_ab_* state increases proportionally with increasing Δ*t*, and continue to be responsive even when the two inducers are separated by 8 hours. There is some expression of GFP in the presence of only inducer **a** (*E_a_*), indicating some basal levels of intB. RFP expression, a proxy for the number of cells in state *S_a_*, begins to increase at *t* = 0 h and drops at time *t* = Δ*t* when inducer **b** is added (Figure S3A). Aligning all of the GFP expression curves by Δ*t* (Figure S4) shows that lower values of Δ*t* not only have lower final GFP expression values, but also have slower rates of GFP production. This is consistent with modeling results because if we assume inducer **b** has an equal probability of entering any one cell, then in case of small Δ*t* (Δ*t* ≤ 4 hours) there is a much larger number of *S_o_* cells and so the rate of *S_a_* → *S_ab_* state conversion will be lower. In the case of Δ*t* > 4 hours, the majority of cells in the population are already in the *S_a_* state configuration, and so the rate of cell state conversion to *S_ab_* will be much higher.

**Figure 4:**
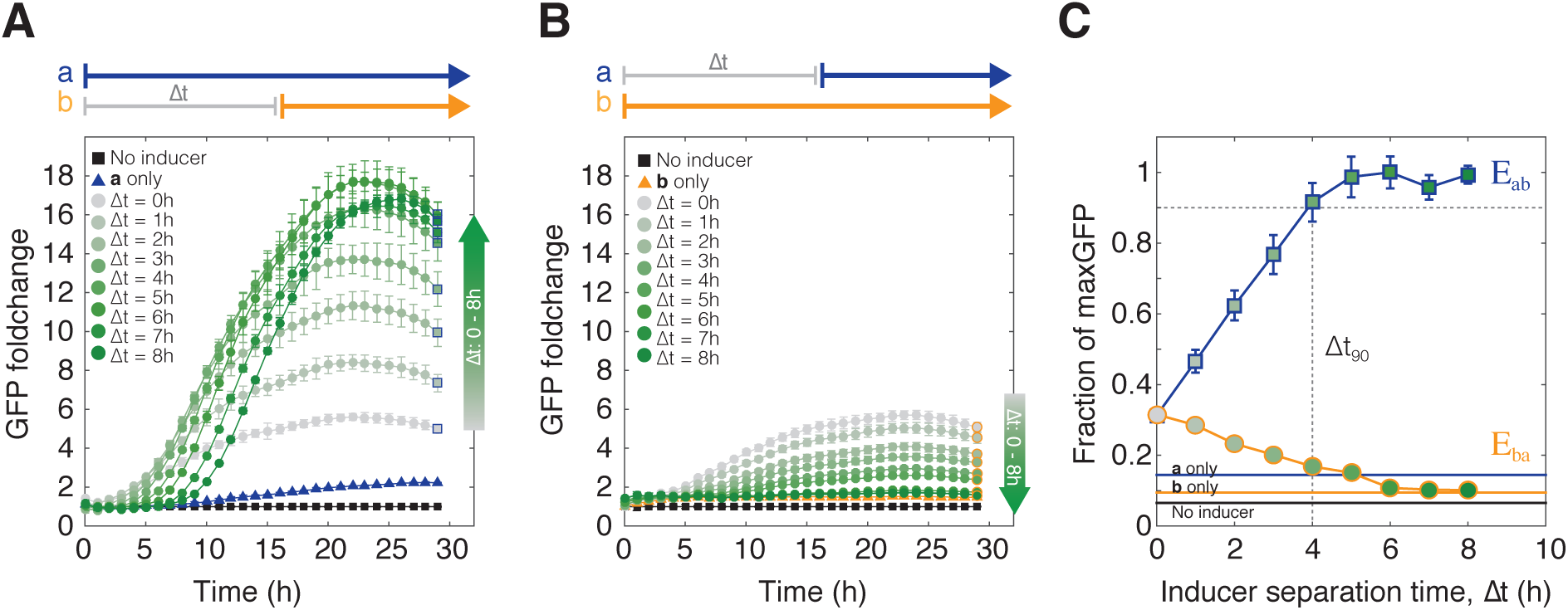
*In vivo* results for varying inducer separation time from Δ*t* = 0 − 8h. A) Populations of cells exposed to an *E_ab_* event sequence. Cell-switching to state *S_ab_* (indicated by GFP fluorescence) begins when inducer **b** (aTc) is added. Maximum normalized GFP fluorescence increases as a function of the inducer separation time Δ*t*. Gray to dark green gradient represents increasing Δ*t* values. Square markers are final endpoint measurements. Error bars represent standard error of the mean. B) Cells exposed to the inverse *E_ba_* sequence of events. GFP fluorescence is inversely proportional to the inducer separation time between **b** and **a**. Circular markers are final endpoint measurements. C) The final fluorescence values for Figure 4 A, B at 30 hours are plotted as a function of Δ*t*. The final fluorescence values have been normalized to the maximum GFP expression. Dotted line marks Δ*t*_90_ detection limit.

When cells are exposed to *E_ba_*, the number of *S_ab_* cells decreases proportionally to Δ*t* (Figure 4B), and there is no RFP expression above background (Figure S3B). In both types of events, the cells maintained their state for up to 30 hours. Raw data for this set of experiments can be found in Figure S5.

We can differentiate between populations that have been exposed to *E_ab_* versus *E_ba_* within one hour of separation time between inducers (Figure 4C). Endpoint GFP fluorescence measurements at 30 hours from Figure 4A,B were normalized by the maximum fluorescence and plotted against Δ*t*. As Δ*t* increases, the GFP-expressing *S_ab_* sub-population increases. The populations that encountered *E_ba_* show decreasing GFP expression as Δ*t* increases, and at Δ*t* = 6 h, the expression is equal to the baseline expression of a ***b*** *only* population, indicating that the addition of inducer **a** after a 6 hour exposure to only inducer **b** has no effect at all. Based on where the GFP fraction exceeds 90% of the max expression, the Δ*t*_90_ detection limit for the experimental system is ~ 4 hours. Finally, the baseline population split when both **a** and **b** are added simultaneously (Δ*t* = 0 h) is not 50% of the maximum GFP expression as expected. This suggests that the integrase flipping rates, *k*_flipA_ and *k*_flipB_, may not be equal and that the basal expression rates, *k*_leakA_ and *k*_leakB_, are non-zero.

### Varying model parameters for integrase activity and basal expression

The parameters for integrase flipping and leaky basal expression were tuned to account for the asymmetrical population responses to *E_ab_* versus *E_ba_* events (Figure 4C). We hypothesized that this asymmetry arises from a combination of unequal integrase activity when searching for and flipping the DNA, and unevenness in protein expression as well as leaky background expression of the integrases (Figure 5). We varied these parameters for intA in the model relative to constant intB parameters. When the relative flipping efficiency parameters of intA (*k*_flipA_) was decreased from 100% of intB efficiency to 0.25-0.75*k*_flipB_, we observed a bias in the baseline population split when Δ*t* = 0 h and both inducers are introduced simultaneously (Figure 5A). Previously in the preliminary model (Figure 3C), the two integrases were assigned equal flipping rates, and the population split was expected to be 50/50. As the flipping rate of intA decreases relative to that of intB, that baseline shifts downwards to favor the more active integrase, intB.

**Figure 5:**
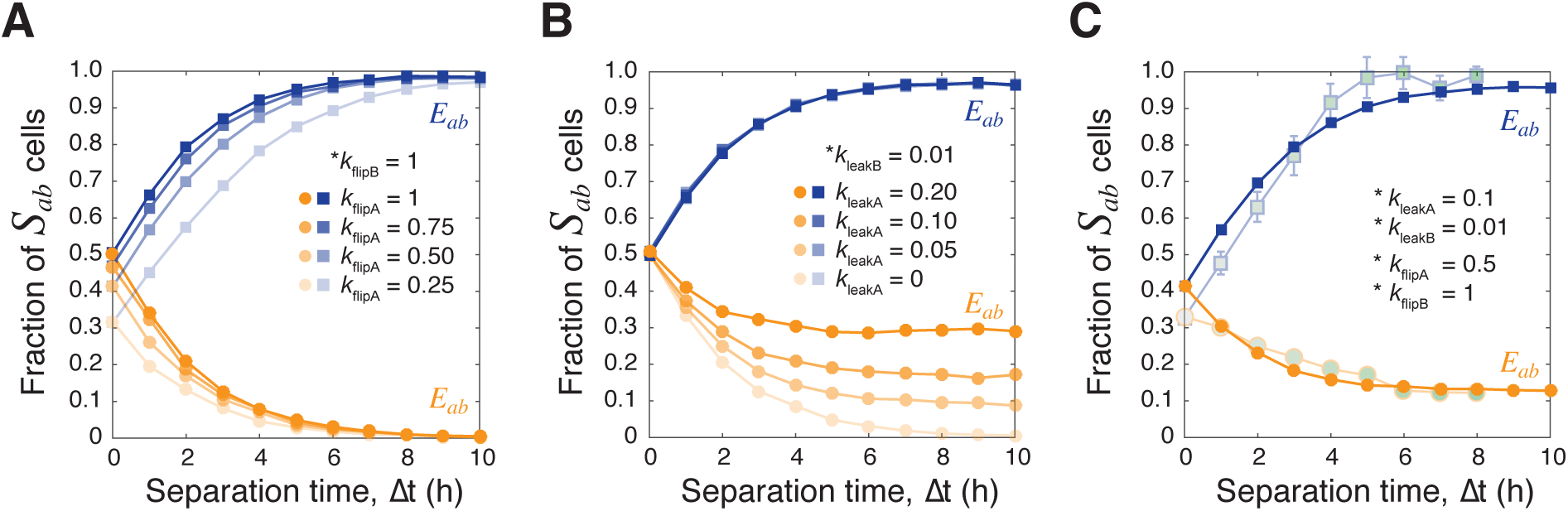
Varying model parameters for integrase flipping and leaky expression. A) As DNA flipping rates of intA (*k*_flipA_) are decreased relative to *k*_flipB_, the population of *S_ab_* cells at Δ*t* = 0 h has a downward shift. Simulations are done with N = 10,000 cells. B) Increasing the leaky expression of intA (*k*_leakA_) changes the minimum number of cells that erroneously switch in to the *S_ab_* state when exposed to *E_ba_*. C) The model was revised to more closely match the experimental data (opaque). The revised parameters are *k*_flipA_ = 0.5hr^−1^, *k*_flipB_ = 1hr^−1^, *k*_leakA_ = 0.1hr^−1^, and *k*_leakB_ = 0.01hr^−1^.

The experimental data show a small population of *S_ab_* cells even when only one of the inducers is present (Figure 4C, ***b*** *only*, ***a*** *only*). When the flipping rates were held equal and the leaky expression of intA (*k*_leakA_) was varied from 0 to 0.2 hr^−1^ (Figure 5B), this increased the baseline minimum number of cells that incorrectly end up in the *S_ab_* state during an *E_ba_* event. The fraction of *S_ab_* cells for Δ*t* >6 h have reached this threshold of minimum leaky expression.

When the combined effects of unequal integrase flipping activity and leaky expression are added to the model (Figure 5C), the separation time Δ*t* versus *S_ab_* population fraction graph generated by the model more closely matches the *in vivo* results. Experimental results (Figure 4C) have been overlaid with reduced opacity to show fit. Based on this qualitative fitting, intA appears to be less active and more leaky than intB, with parameters changed to *k*_flipA_ = 0.5hr^−1^ and *k*_leakA_ = 0.1hr^−1^. This suggests that to the high leaky expression of intA, around 10% of the population will “detect” *E_ab_* and be in state *S_ab_* even when no inducer **a** has been introduced. Although intB also has some basal level of expression, it is more difficult to ascertain the percentage of cells that erroneously switch to state *S_b_* since *S_b_* does not have a fluorescent output.

Finally, the Δ*t*_90_ detection limit can be tuned by increasing or decreasing the overall production rate *k*_prod*_ (* = A or B) (Figure S6). In this particular implementation of the temporal logic gate, Δ*t*_90_ is ~ 4 hours. Within this 0 to 4 hour window, the *S_ab_* population fraction can be used to uniquely determine Δ*t*. Outside of this time window, the only assertion that can be made is that Δ*t >* 5 hours. If *k*_prod*_ (* = A or B) were higher, the integrases would accumulate faster in each cell and increase the probability of DNA flipping. Such a system would have a lower Δ*t*_90_ but also higher resolution of events within the narrower *t* = 0 to *t* =Δ*t*_90_ hour time window. This is because with higher protein production rates, cells commit to a final state faster, thus shrinking the effective window of time in which events could be resolved. However, within that smaller time window, *S_ab_* populations fractions would also be measurably different at much smaller intervals, and so Δ*t* could be resolved with much higher resolution. On the other end of the spectrum, if protein production were slow, the stochastic DNA recombination events happen less frequently, resulting in a population that is more sensitive to inputs for a longer period of time, but would have lower resolution overall since the population fractions are not changing as quickly.

### Deducing inducer pulse width

Using the fraction of GFP-expressing *S_ab_* cells alone, we can determine Δ*t* values up to a Δ*t*_90_ limit for any given sequence of two step inputs. Now consider a pulse type of event, in which inducer **a** begins at time *t* = 0 h and remains constant throughout and inducer **b** is introduced as a finite pulse at time *t* = Δ*t* h (Figure 6A). The start time of inducer **a** then becomes a reference for when the entire system is activated and ready to detect inducer **b**. Cell states are measured via fluorescence at time *t*_end_, where *t*_end_ > 24 hours. Modeling results presented in this section are using the refined set of parameters defined in Figure 5C and Table S5.

**Figure 6:**
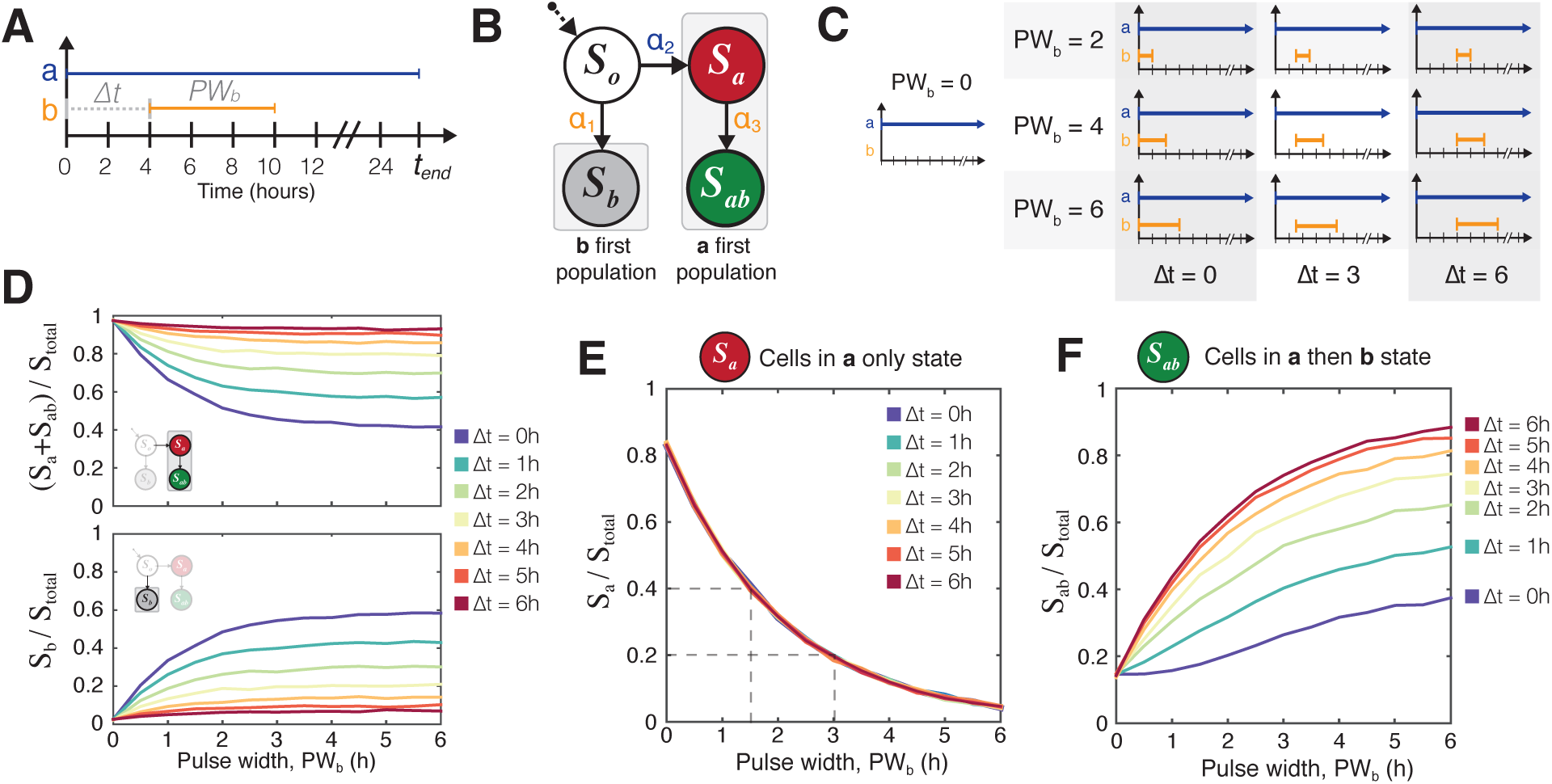
Simulation results for pulse width modulation. Simulations were done with revised parameters found in Figure 5C. A) Inducer **a** can be used as a reference signal against which to measure the time and duration of the inducer **b** pulse. B) The population eventually divides into one of two partitions: those that see inducer **a** first and those that see inducer **b** first. Only if a cell has entered the *a first* pathway does it have the possibility to express RFP or GFP. Furthermore, *S_a_* can be thought of as a necessary precursor to *S_ab_*. C) A matrix illustrating a subset of the Δ*t* and values to be tested. D) Simulation results show that for any given Δ*t*, the number of cells in *S_b_* = total number of cells − (*S_a_* + *S_ab_*) E) The fraction of the population in the *S_a_* state is totally independent of Δ*t* and depends only on the pulse duration of inducer **b**. F) Once is known, then the fraction of the population in *S_ab_* state can be used to find the time at which the pulse of inducer **b** began.

If either of the two inducers is present in the media to some limit *t*_end_, we would expect all of the *S_o_* cells will end up in one of two populations (Figure 6B). Cells that encounter inducer **b** first will be in the *S_b_* state, while cells that encounter **a** first will either be in the *S_a_* or *S_ab_* states. In the previous sections, once an inducer was added to the population, it was not removed, and the assumption was made that at times greater than 24 hours, only a negligible number of *S_o_* cells remained. This type of step function induction also meant that only the number of *S_ab_* cells (GFP) was needed to uniquely determine the separation time Δ*t* because *any and all* cells that had switched to *S_a_* would eventually become *S_ab_*.

However, in the case of a transient pulse, some cells that are in the *S_a_* state (RFP) will not ever encounter inducer **b**. Assuming the *k*_leakB_ is small, these cells will remain in the *S_a_* state. Therefore, the population of ***a*** *first* cells equals *S_a_* + *S_ab_*. We simulated a matrix of populations exposed to varying inducer separation times (Δ*t*) and inducer **b** pulse widths (*PW_b_*) to measure the resolution of detectable events (Figure 6C). In simulation (Figure 6D), we can see that the two populations mirror each other to add up to 100% of the total cells (N = 10,000 cells).

Since the step induction of **b** is equivalent as a pulse of infinite length (*PW_b_* = ∞), and no cells remain in state *S_a_* when *PW_b_* = ∞, then perhaps the number of *S_a_* cells can be used to deduce information about pulse width. In *silico*, we can test this hypothesis by running a matrix of simulations with varying Δ*t* and *PW_b_*. In Figure 6E, we see that the fraction of *S_a_* cells over the total number of cells decreases monotonically with increasing *PW_b_*. Due to some non-zero *k*_leakB_, the curves produced by different Δ*t* values do not completely overlap but the different is indistinguishably small. The maximum number of *S_a_* cells does not go to 1 at *PW_b_* = 0h because of leaky intB expression (*k*_leakB_ = 0.01).

If the fraction of *S_a_* cells depends only on the duration of *PW_b_*, the fraction of RFP cells relative to the total number of cells can be used to uniquely determine the pulse length of inducer **b**, *PW_b_* (Figure 6E). Once *PW_b_* is known, the fraction of GFP-expressing cells (*S_ab_*) can be used to uniquely determine the time between inducers, Δ*t* (Figure 6F). Furthermore, the genetically encoded state means that these population fractions should be maintained and measurable at a time, *t_end_*, that is much later than the time of the events.

These conclusions can be extended in simulation to create a scatterplot of *S_a_* cells versus *S_ab_* cells in a population (Figure 7A) over an 11×11 parameter matrix varying Δ*t* and *PW_b_* from 0 – 6 hours in increments of 0.5 hours (Additional plots in Fig. S7). Each point on the chart in Figure 7A represents a simulated population (N = 10,000) exposed to a unique combination of Δ*t* and *PW_b_* values. Vertical lines represent the *sa*me *PW_b_* value, and points with the *sa*me shape and color have the *sa*me Δ*t* value. (See also Figure S7, Table S2). The simulation results suggest sufficient resolution of events as long as *PW_b_* and Δ*t* values are between 0 to 4 hours. For any single value of *PW_b_*, we can follow the increasing Δ*t* values vertically and see that the population response *sa*turates after 4.5 hours resulting in overlapping between populations with 4.5 < Δ*t* < 6 hours. We can trace any individual Δ*t* value horizontally from right to left, and observe that the points begin to cluster and overlap when 4.5 < *PW_b_* < 6 hours. These simulation data suggest that there should be some defined detection range of Δ*t* and *PW_b_* where each possible combination of the two is uniquely identifiable.

**Figure 7:**
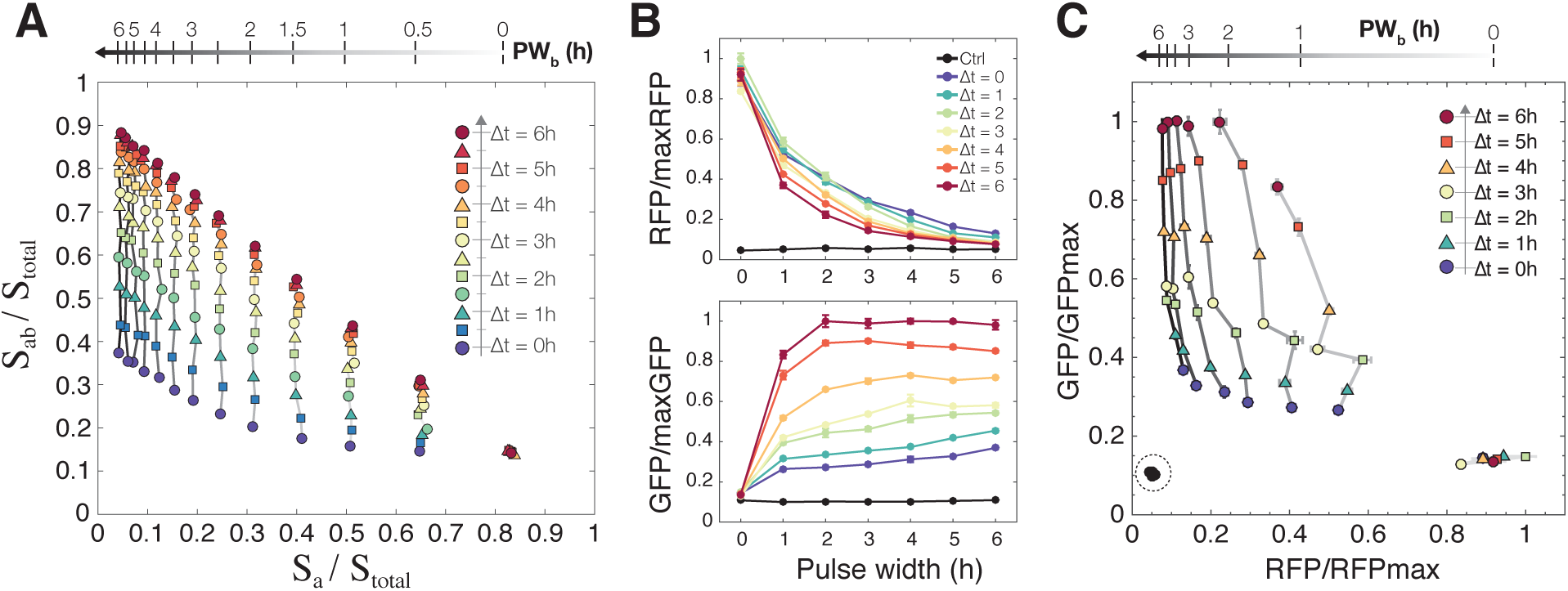
Determining arrival time and pulse duration of inducer **b** with population fractions. A) Simulation results from testing an 11 × 11 matrix of parameters with Δ*t* and *PW_b_* varying from 0 – 6 hours in increments of 0.5 hours. Each point represents a population of 10,000 cells. Increasing *PW_b_* goes from right to left, and increasing Δ*t* goes from bottom to top. B) Experimental results showing RFP and GFP expression as a function of increasing Δ*t* and *PW_b_*. Fluorescence values have been normalized to the highest GFP and RFP fluorescence in the *sa*mple set. Experimental results from exposing temporal logic gate *E. coli* populations to varying *PW_b_* and Δ*t* values (0 – 6 hours, 0.01%/vol L-ara, 200 ng/ml aTc, measurements taken at 48 hours). C) A scatterplot of each population using their RFP and GFP expression as coordinates. The non-induced control *sa*mples are circled on the bottom left, and the *sa*mples with *PW_b_* = 0h are on the bottom right. *Sa*mples with the *sa*me *PW_b_* are connected with a solid line, and line darkness represents increasing *PW_b_* duration. *Sa*mples with the *sa*me Δ*t* are shown with the *sa*me colored shape marker and increasing Δ*t* goes from bottom to top.

Experimentally, we tested a 7 × 7 matrix of varying Δ*t* and *PW_b_* (0 – 6 hours, 1 hour increments) on independent populations of the temporal logic gate *E. coli* strain (Figure 7B). All populations, except for the control, were exposed to inducer **a** (L-ara 0.01%/vol) at time *t*_0_ to *t*_end_. Pulses of inducer **b** (aTc, 200ng/ml) were achieved by *sa*mpling 5ul of the population and diluting 1:100 into fresh media with only inducer **a** (M9CA + 0.01%/vol L-ara). Fluorescence measurements were taken at 24 and 48 hours. For all values of Δ*t*, the number of *S_a_* cells (RFP) is highest when there is no exposure to inducer **b** (*PW_b_* = 0h) and decreases monotonically as a function of *PW_b_* (Figure 7B, top). RFP and GFP fluorescence expression has been normalized by the highest RPF and GFP expression values in the *sa*mple set. Our numerical simulations predicted a complete overlap of the *S_a_* curves (Figure 6E), and the experimental results are consistent with those predictions though there is some downwards drift with increasing Δ*t*. We see a more pronounced separation of the Δ*t* curves when we look at GFP expression, a representation of the number of cells in the *S_ab_* state. The number of *S_ab_* cells is dependent on both Δ*t* and *PW_b_* and increases proportionally with both increasing **b** pulse duration and inducer separation time.

Even with these bulk fluorescence measurements, we can resolve the different populations that result from varying Δ*t* and *PW_b_* values (Figure 7C). As with Figure 7A, each point on the graph represents an independent population of cells (OD ~ 0.7). For any one value of Δ*t*, increasing *PW_b_* is inversely proportional to RFP expression, or *S_a_* state cells. For any one value of *PW_b_*, the RFP expression remains relatively constant with increasing Δ*t*, while GFP, or the number of *S_a_*_b_ state cells, increases 3-fold from Δ*t* = 0 to 6 hours. In the case where there is **b** pulse (*PW_b_* = 0h), the amount of RFP fluorescence increases with *sa*mpling time Δ*t*, but there is negligible expression of GFP. Populations with different *PW_b_* exposures are well separated up to 4 hours though there is some drift in RFP as Δ*t* increases. All of the populations exposed to either or both of the inducers are out of the range of the no inducer controls (indicated by dotted circle). An additional table of this data sorted by fluorescence expression can be found in Table S2.

This method of profiling is only valid if the fraction of *S_a_* state cells can be used as a measure of *PW_b_* that is independent of Δ*t*. To ensure that this is not an artifact of simulation or experimental systems, we also mathematically analyzed the equation (1) for ℙ*_t_*(*S_a_*), which represents the fraction of cells with DNA state *S_a_* at time *t*.

If inducer **a** is used as a constant reference signal, all cells transition into either of either of *S_a_, S_b_* or *S_ab_* state, thus ℙ_∞_(*S_a_*) = 1 − (ℙ_∞_(*S_b_*) + ℙ_∞_(*S_ab_*)). If we assume that the basal leaky expression of intB is zero (*k*_leakB_ = 0), ℙ*_t_*(*S_b_*) + ℙ*_t_*(*S_ab_*) = 0 holds for *t* ≤ Δ*t*, since there is no intB that turns DNA state into *S_b_* or *S_ab_*. Then, we can show that ℙ*_t_*(*S_b_*) + ℙ*_t_*(*S_ab_*) is dependent only on *PW_b_*, the duration of the pulse width of inducer B, for *t* > Δ*t* (see SI for details). Thus we can conclude that ℙ*_t_*(*S_a_*) = 1 − (ℙ*_t_*(*S_b_*) + ℙ*_t_*(*S_ab_*)) is dependent on *PW_b_* but not on Δ*t*, implying that the number of cells in the *S_a_* state is a function of *PW_b_* as t → ∞ (See SI for derivation).

### Practical use and calibration

Curve-fitting methods were used to automatically convert experimentally measured RFP and GFP fluorescence into *PW_b_* and Δ*t* values and to evaluate the resolution with which population fluorescence ratios can be used to determine inducer separation time and pulse duration. Using the experimental data from Figure 7B,C, we generated fitting curves for *PW_b_* as a function of RFP/maxRFP(R), and for Δ*t* as a function of both GFP/maxGFP(*G*) and *PW_b_* (Figures S8 – S9, Table S1). We will denote these functions with *PW_b_*(*R*) and Δ*t*(*G, PW_b_*), respectively. The functions *PW_b_*(*R*) and Δ*t*(*G, PW_b_*) can then be used to generate a mesh of predicted *PW_b_* and Δ*t* values for any given normalized fluorescence values (Figure 8A, equations in SI).

**Figure 8:**
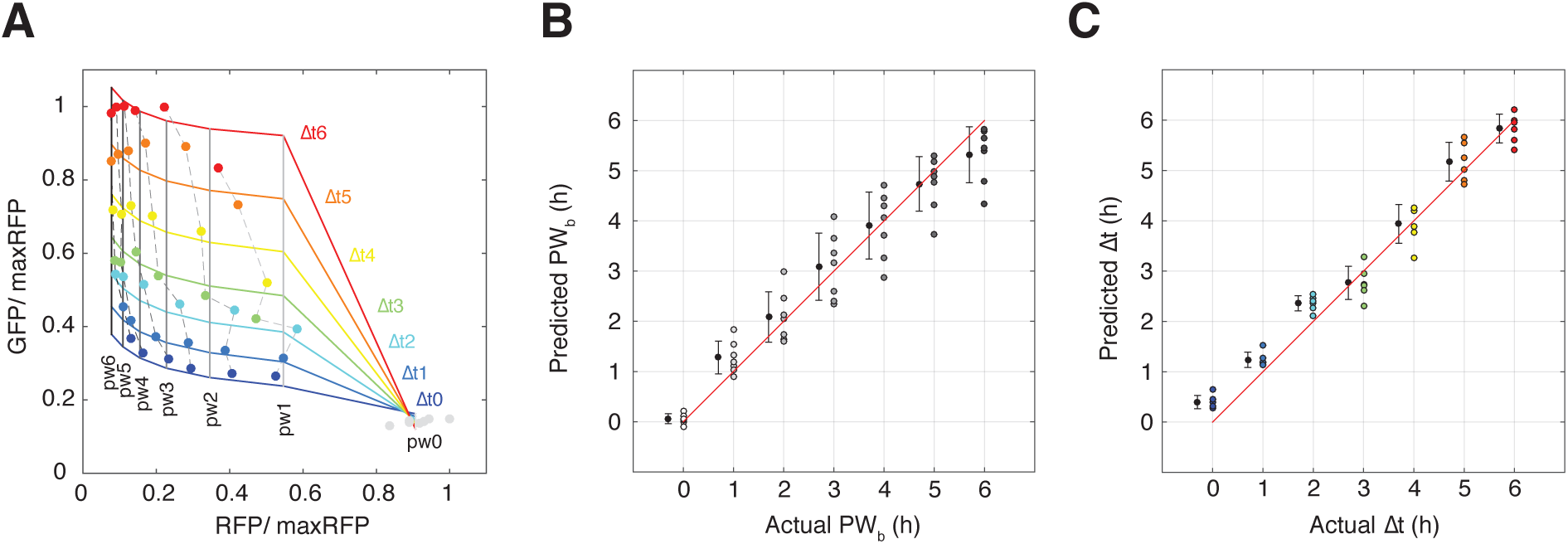
Determining prediction resolution for *PW_b_* and Δ*t* from fluorescence data. A) A mesh generated from fitted curves for *PW_b_* as a function of RFP/maxRFP(R) and Δ*t* as a function of pulse width and GFP/maxGFP(G). Experimental data is overlaid. B) Comparison of actual versus predicted *PW_b_* values generated by fitted function *PW_b_*(*R*). For each actual *PW_b_* value, the average of the predicted *PW_b_* values with ± 1 standard deviation (Slightly offset on the *x*-axis for better comparison). C) Comparison of actual versus predicted Δ*t* generated by the fitted function Δ*t*(*G, PW_b_*). For each actual Δ*t* values, the average of the predicted Δ*t* with ± 1 standard deviation (Slightly offset on the *x*-axis for better comparison).

The predicted values were compared against the actual values to determine the approximate time window with which a specific *PW_b_* or Δ*t* can be resolved. For each actual value of *PW_b_* and Δ*t*, we calculated the average and standard deviation for the set of predicted values. The standard deviation allows us to visualize the range for which the majority of predictions will fall for any given actual value. For instance, a *PW_b_* of 0 hours can be detected ±0.25 hours, but as *PW_b_* increases, this prediction window widens and for *PW_b_* ≥ 3 hours, the resolution of detection is closer to ±1 hour (Figure 8B). Similarly, predicted values of Δ*t* fall within ±0.25 hours for 0 < Δ*t* < 3 hours and increase to ±0.5 hours when Δ*t* ≥ 3 hours (Figure 8C). Using these fitting functions, we can also pre-generate a reference table that converts normalized RFP and GFP fluorescence data into predicted *PW_b_* and Δ*t* values (Table S3).

## Discussion

We have designed and implemented a temporal logic gate that takes advantage of the population dynamics to collectively sense and record sequences of transient chemical inputs. As with all engineered systems, proper calibration of these temporal logic gate populations will be required prior to deployment in the “field.” We envision a process similar to the one described in this report. First, experimental populations are exposed to a matrix of *PW_b_* and Δ*t* values. This will set the maximum and minimum fluorescence for RFP and GFP and provide necessary data for determining the Δ*t*_90_ limit and producing the fitting functions *PW_b_*(*R*) and Δ*t*(*G, PW_b_*). Once the fitting functions have been determined, values for *PW_b_* and Δ*t* for experimental samples can be estimated within ± 0.25 to 1 hour of the actual values. A calibrated table could also be generated and used for as a reference for samples that have been exposed to unknown conditions.

The stochastic nature of molecular processes often presents a significant barrier to homogenous outputs from an engineered population of cells. This implementation of event detection via population fractions takes advantage of stochastic and heterogenous individual responses to environmental conditions in order to map final population fractions back to unique sequences and durations of chemical events. The sensitivity of the system and the Δ*t*_90_ detection limit could be modulated by increasing or decreasing protein production rates via tuning of plasmid copy numbers, signal concentration, or transcription/translation sequences. The use of digital cellular outputs combined with the analog population response creates event detection systems that are more robust to stochasticity and can be tuned more easily.

As a proof-of-concept, we have used the common laboratory inducers L-arabinose and aTc as inputs, but we hope that our temporal logic gate system can be used modularly with any biosensors of choice. In particular, we believe there are possibilities for detection of miRNAs and biofilm formation. Stable populations of microRNAs (miRNAs) circulating in the blood have generated a lot of interest as biomarkers for human health (Cortez *et al*, 2011). These short (~ 20-30nt) regulatory RNAs have been shown to have sequential tissue-specific expression signatures that correlate with pregnancy, tumor formation, and other diseases (Gilad *et al*, 2008; Mitchell *et al*, 2008), and synthetic biology has developed many customizable RNA sensors (Friedland *et al*, 2009; Green *et al*, 2014). Another possible application of this would be detection of harmful biofilms. Biofilms are self-assembling, highly structured, multi-species consortia that develop in stages and have sophisticated networks of interaction and function (Stoodley *et al*, 2002; Flemming and Wingender, 2010; Elias and Banin, 2012). Unnatural biofilm development in environments such as industrial water sources or waste streams can be both harmful for both the natural environment and the industrial mechanisms. Detection of biomarkers for known strains of biofilm colonizers would provide early warning of changing ecosystems, and although we do not yet fully understand these networks, it is known that quorum-sensing plays a critical role in the process. Quorum-sensing molecules and receptors are available in the synthetic biology toolbox and so may provide an accessible way of detecting the sequential colonization of different microbes. Field deployment of engineered bacteria will likely involve transient signals, low-nutrient environments, and possibly even other microbial competitors (i.e. soil, flowing rivers, the digestive tract). We used minimal media in this study to better approximate low-nutrient environments, and anticipate further characterization in more customized ‘local’ environments (i.e. gut model or air model or soil model) and with hardier microbial chassis.

Finally, this study focused on the population outputs as indicators of past events, but we believe that this temporal logic gate could be used to reliably differentiate a single strain into controlled sub-populations via input pulse order, duration, and frequency. In recent years, it has been recognized that many natural systems modulate cellular behavior not only by changing the concentration of signaling molecules but also by regulating signal pulse frequency (Cai *et al*, 2008; Lin *et al*, 2015). If we consider the fluorescent proteins GFP and RFP in this circuit as simply placeholders for downstream genes, then this system could easily be applied as a top-down population differentiator. By modulating the sequence of inputs, one could systematically predict and create mixed populations of genetically differentiated cells. As the scientific community turns towards further understanding of microbiomes and multi-cellular consortia, engineered bacteria populations could be used not only as a tool for investigating the activities of natural communities but also as a way to build synthetic communities from the ground up.

## Materials and methods

### Cell strains and plasmids

All plasmids used in this study were designed in Geneious 7.1 (Biomatters, Ltd.) and made using standard Gibson isothermal cloning techniques. Integrases Bxb1 and TP901-1 are on a high-copy plasmid (pVHed05, plasmid map in Figure S9) with a ColE1 origin of replication (original template from the Dual Recombinase controller (Bonnet *et al*, 2013), Addgene Plasmid 44456). Integrase A (Bxb1) is behind a Ptet promoter and integrase B (TP901-1) is behind a PBAD promoter. The plasmid has been modified with an additional TetR gene. The temporal logic gate was integrated into the Phi80 site on the *E. coli* chromosome using CRIM integration (Haldimann and Wanner, 2001) and screened for single integrant colonies. The integration plasmid template and DH5*α*-Z1 strain were generously provided by J. Bonnet and D. Endy and modified to contain the temporal logic gate (pVHed07, plasmid map in Figure S9).

Additional DNA and oligonucleotides primers were ordered from Integrated DNA Technologies (IDT, Coralville, Iowa).

A custom formulation of M9CA media was used for all experiments. The media contained 1× M9 salts (Teknova, M1906) augmented with 100mM NH_4_CL, 2mM MGSO_4_, 0.01% casamino acids, 0.15 *μ*g/mL biotin, 1.5 *μ*M thiamine, and 0.2% glycerol, and then sterile filtered (0.2 *μ*m).

### Simulations of the model

The stochastic simulation algorithm by Gillespie (Gillespie, 1977) was implemented to generate the sample paths of individual cells using the Markov model (see Table S6 for the definitions of Markov transitions and transition rates). All simulation runs and their analyses were done with MATLAB (R2014b,The MathWorks, Inc.). All simulated populations were done with 10,000 individual cell trajectories.

### Experimental methods

Prior to all experiments, cells were grown overnight from plate cultures in M9CA for two days, then diluted to OD 0.1 and recovered for 4-6 hours at 37*^o^*C. L-arabinose and anhydrous tetracycline (aTc) were used as inducers **a** and **b**, respectively. L-ara was used a concentration of 0.01% by volume, and aTc was used a concentration of 200 ng/ml (450nM). All media contained the antibiotics chloramphenicol (source and concentration) and kanamycin (source and concentration). All experiments were performed with the aid of timed liquid handling by a Hamilton STARlet Liquid Handling Robot (Hamilton Company).

For step function experiments, the cells were diluted to OD 0.1 into a 96-well matriplate (Brooks Automation, Inc., MGB096-1-2-LG-L) with 500*μ*l total volume in M9CA. Cultures were incubated at 37^o^C in a BioTek Synergy H1F plate reader (BioTek Instruments, Inc.) and inducers were added at appropriate time by the Hamilton robot. OD and fluorescence measurements (superfolder-GFP ex488/em520, mKate2-RFP ex580/em610) were taken by the BioTek every 10 minutes.

For the pulse experiments, pulses were achieved through dilution of the culture into fresh M9CA media containing 0.01% L-arabinose. 5 *μ*l of the culture was sampled and diluted into 500 *μ*l of fresh M9CA + 0.01% L-ara to achieve pulsatile exposure to aTc. 96-well deep-well plates containing the diluted cultures were then incubated at 37^o^C and fluorescence measurements were taken at 24 and 48 hours in the plate reader.

Analysis of experimental data was done using custom MATLAB scripts. All depicted error bars are standard error of the mean. Fitting of curves was done in MATLAB.

## Acknowledgements

The authors would like to thank J. Bonnet and D. Endy for the initial plasmids used in this work, S. Sanchez for critical assistance in automation and liquid handling, and C. Hayes for discussions.

V.H. is supported by the U.S. Department of Defense (DoD) through the National Defense Science & Engineering Graduate Fellowship (NDSEG) Program. Y.H. is supported by JSPS Fellowship for Research Abroad. Research supported in part by the Institute for Collaborative Biotechnologies through grant W911NF-09-0001 from the U.S. Army Research Office. The content of the information does not necessarily reflect the position or the policy of the Government, and no official endorsement should be inferred.

## Author Contributions

V.H. conceived of the circuit design, constructed the necessary experimental strains, performed experimental work, and ran model simulations. Y.H. developed the stochastic model and derived the mathematical results. P.W.K.R provided feedback and guidance on data analysis and interpretation. R.M.M provided feedback and guidance on overall project vision, circuit design, and interpretation of results.

